# Generation of specialized blood vessels through transdifferentiation of lymphatic endothelial cells

**DOI:** 10.1101/2021.04.28.441726

**Authors:** Rudra N. Das, Ivan Bassi, Yanchao Han, Giuseppina Lambiase, Yaara Tevet, Noga Moshe, Stav Safriel, Julian Nicenboim, Matthias Brückner, Dana Hirsch, Raya Eilam-Altstadter, Wiebke Herzog, Kenneth D. Poss, Karina Yaniv

## Abstract

The lineage and developmental trajectory of a cell are key determinants of cellular identity. Yet, the functional relevance of deriving a specific cell type from ontologically distinct progenitors, remains an open question. In the case of the vascular system, blood and lymphatic vessels are composed of endothelial cells (ECs) that differentiate and diversify to cater the different physiological demands of each organ. While lymphatic vessels have been shown to originate from multiple cell sources, lymphatic ECs (LECs) themselves seem to have a unipotent cell fate. In this work we uncover a novel mechanism of blood vessel formation through transdifferentiation of LECs. Using advanced long-term reiterative imaging and lineage-tracing of ECs in zebrafish, from embryonic development through adulthood, we reveal a hitherto unknown process of LEC-to-BEC transdifferentiation, underlying vascularization of the anal fin (AF). Moreover, we demonstrate distinct functional implications for deriving AF vessels from either LECs or BECs, uncovering for the first time a clear link between cell ontogeny and functionality. Molecularly, we identify Sox17 as a negative regulator of lymphatic fate specification, whose specific expression in AF LECs suppresses its lymphatic cell fate. Finally, we show that akin to the developmental process, during adult AF regeneration the vasculature is re-derived from lymphatics, demonstrating that LECs in the mature fish retain both potency and plasticity for generating specialized blood vessels. Overall, our work highlights a novel mechanism of blood vessel formation through LEC trans-differentiation, and provides the first *in vivo* evidence for a link between cell ontogeny and functionality in ECs.

---

After the emergence of lymphatic endothelial cells (LECs) in the early embryo^1^, lymphatic vessels undergo various organ-specific specializations^2^, the knowledge of which is restricted to a few studies. In zebrafish, few instances of adult-specific lymphatic diversification were described to take place during metamorphosis^3^, i.e. the larva-to-juvenile transition^4,5^. Here, we focus on the anal fin (AF), an adult-specific structure, first detected when the larva reaches ∼5.2 mm in length (∼14-16 dpf) (Fig. 1a), whose formation and maturation can be tracked *in vivo* due to its peripheral location and small tissue depth. Zebrafish possess 3 unpaired fins (dorsal, caudal and anal), each composed of cartilaginous (endo-skeletal) and dermal (exo-skeletal) bones^6,7^, with the latter receiving vascularization through a well-organized system of veins and arteries^8,9^ (Fig. 1a’-a’’). In contrast, the status of lymphatic vessels in these organs remains largely unknown.

**Fig. 1:**
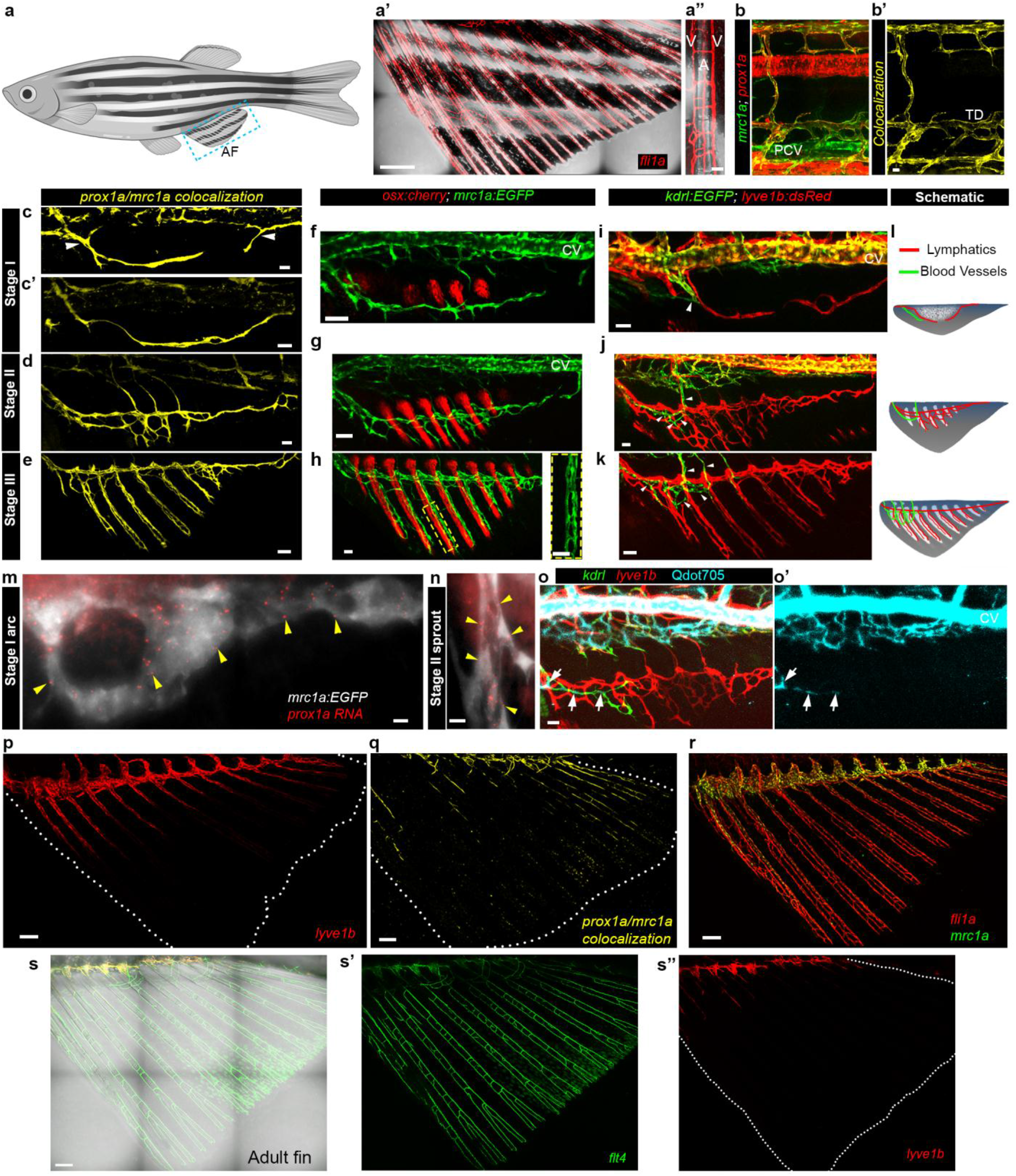
Stepwise formation of the juvenile anal fin vasculature reveals a unique dynamics of lymphatic marker expression. a, Illustration of an adult zebrafish indicating the AF location (blue box). a’-a’’, Mature vascular network in adult (3 mpf) AF (a’) and in a single fin ray (a’’) of *Tg(fli1a:dsRed) fish*. Veins (V) and arteries (A) are annotated in a’’. b-b’, Lymphatic labeling in the trunk of *Tg(mrc1a:EGFP)* and *Tg(prox1a:RFP)* larvae at 14dpf; co-localization channel in b’ is depicted in yellow. c-e, *prox1a/mrc1a* colocalization channel showing lymphatics in stages I,II,III of AF development. Arrowheads in (c) point to anterior and posterior sprouts. f-h, Osteoblast (*osx:cherry*) and i-k, blood vessel (*kdrl:EGFP*) development shown for the three stages of AF development. Lymphatics are labeled by *mrc1a:GFP* in f-h, and by *lyve1:dsRed* in i-k. Inset in h shows a lymphatic vessel growing through a single ray. Arrowheads in i,j,k denote the restricted dorso-rostral position of the blood vasculature. l, Schematic illustrating the growth of lymphatic and blood vessels along with the bones of the developing AF through stages I-III. m-n, smFISH images showing *prox1a* RNA (yellow arrowheads) in *mrc1a*+ vessels of stage I (m) and II (n) AFs. o-o’, Distribution of Qdot705 in stage II AF, following intravascular injection. Arrows point to Qdot705 labeled-*kdrl+* blood vessels within the AF. Lymphatics are labelled by *lyve1b:dsRed*. p-q, Lymphatic markers are lost in stage IV AFs (demarcated by white dots), as depicted in *lyve1:dsRed* (p), and *prox1a/mrc1a* (q) juvenile fish. r, All vessels are present in stage IV AFs as shown by *fli1a:dsRed*, along with lymphatic *mrc1a*. s-s’’, Differential expression of *flt4:mCitrine* (s’) and *lyve1b:dsRed* (s’’) in the adult AF (3mpf). Scale bars, 500µm (a’), 150µm (q); 100µm (p,r); 50μm (a’’,e,f,g,k), 30μm (c,c’,d,h,i,j,o), 20µm (b’), 5µm (n), 3µm (m). PCV, Posterior cardinal vein; CV, Cardinal vein; TD, Thoracic duct.

To study the diversification of LECs during metamorphosis, we utilized double transgenic zebrafish-*Tg(mrc1a:egfp)*^4^ and *Tg(prox1aBAC:KalTA4-4xUAS-E1b:uncTagRFP)*^10^, whose co-localization reliably labels *bonafide* lymphatic vessels (Fig. 1b,b’ and Extended Data Fig. 1a-d’) that are not connected to blood circulation (Extended Data Fig. 1e,e’). Recurrent confocal imaging (every 24-72 hrs) of individual animals between 15-30 dpf allowed us to define and characterize four stages of AF formation based on morphological changes in the bone structures (Extended Data Fig. 1f-i; Table 1), along with establishment of its vascular component. Prior to the development of the AF, the median fin fold area is completely devoid of endothelial cells (ECs) (Extended Data Fig. 1j,j’). Later on, as AF development is initiated by mesenchyme condensation^11,12^ (Extended Data Fig. 1f), lymphatic sprouts arising from the thoracic duct (TD) and the cardinal collateral lymphatic vessel (CCL)^4^, enter first the anterior, and then the posterior poles of the AF (Fig. 1c, arrowheads, Extended Data Fig. 2a). These sprouts merge to form a lymphatic arc at the distal margin of the bulging mesenchyme (Fig. 1c’, Extended Data Fig. 2a’). As the fin rays appear (Fig. 1f) and start elongating ventrally, lymphatic vessels begin ramifying into a plexus (Fig. 1d, g, Extended Data Fig. 2b) that sends sprouts which grow along the rays (Fig 1e,h, Extended Data Fig. 2c). Along with the appearance of the initial lymphatic sprouts, *Tg(kdrl:EGFP)*^13^ labelled blood vessels also penetrate the fin fold (Fig. 1i). However, unlike the lymphatic sprouts, they remain restricted to the dorso-rostral area, close to the fin-trunk junction (Fig. 1j,k, arrowheads). The notable prevalence of lymphatic over blood vessels in the developing fin (Fig. 1l) is surprising, given that for the most, organogenesis requires active blood vascularization that supplies oxygenated blood and nutrients to support growth. Lymphatic formation, in contrast, is known to lag behind the blood vasculature in most organs^2,5,14^.

**Fig. 2:**
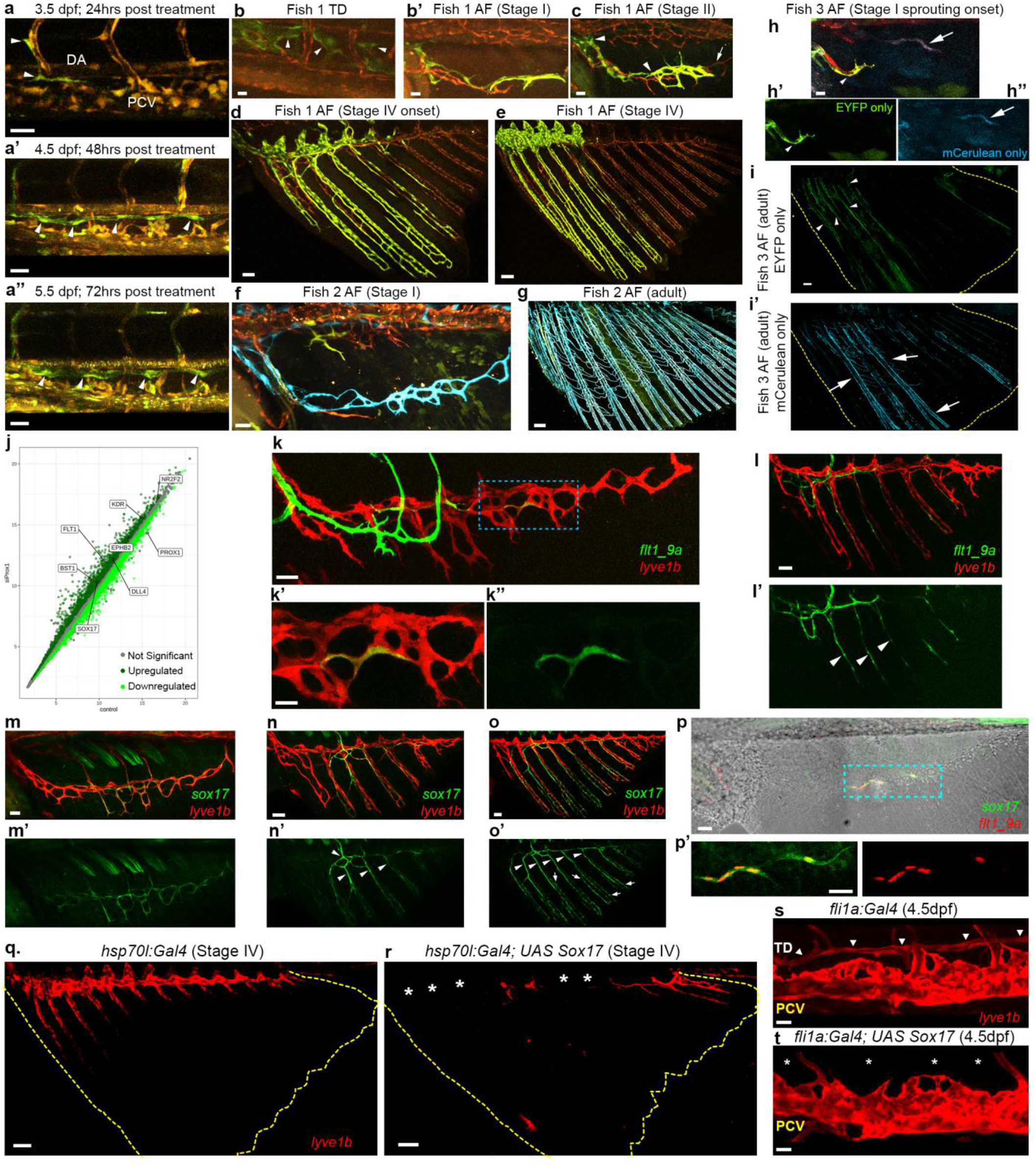
LECs generate BECs through a *sox17*-controlled program. a-a’’, Confocal images of *flibow larvae* showing distinct labeling of lymphatic components in the trunk starting from 24hrs after Cre induction. a, ECs in the early TD of 3.5 dpf embryo labelled in green (arrowheads), are detected 24hrs after heat and 4-HT treatments. a’-a’’, Reiterative imaging of the same embryo at 4.5dpf (a’) and 5.5dpf (a’’) shows the growing TD composed of green ECs (arrowheads). b-e, Reiterative confocal imaging of the same *flibow* fish (fish 1) showing generation of AF vessels from trunk lymphatics. Green ECs, specifically detected in the TD (b, arrowheads) and AF (b’) at stage I, proliferate to occupy the anterior portion of the AF as seen in stages II (c) and IV (d,e). The posterior sprout was not labelled (non-recombinant red, arrow), hence the posterior part of the AF remained unlabeled (d,e). f-g, Confocal images of *flibow* fish (fish 2) showing the final mature AF vasculature to be derived from early AF lymphatic sprout. Stage I lymphatic arc was generated by distinctly labelled blue ECs (f), that completely vascularized the AF of the adult fish (3mpf) (g). h-i’, Confocal images of *flibow* fish (fish 3) showing ECs from the initial blood vessel sprout (h,h’, EYFP, arrowhead) occupying a restricted dorso-rostral position in the mature AF (demarcated by yellow dashed lines) (i), unlike the lymphatic sprout (h,h’’, mCerulean, arrows) which span the entire fin (i’). j, Scatterplot showing rlog expression of genes, in PROX1 suppressed HDLECs, that are upregulated (dark green), downregulated (light green) or did not change significantly (grey). Genes of interest are annotated in the plot. k-k’’, *flt1_9a:EGFP* expression is detected in individual *lyve1:dsRed*-positive LECs (blue dashed ROI) of the early AF lymphatic plexus. k’ and k’’ show high magnification of separate channels. l-l’, *flt1_9a:EGFP* is expressed in the prospective arteries of stage III AF (arrowheads). m-o’, Expression of *sox17*^*EGFP*^ along with *lyve1:dsRed+* lymphatics at stages II (m-m’), III (n-n’) and IV (o-o’) of AF growth. Expression in presumptive arteries (arrowheads) and veins (arrows) is shown. p-p’, *sox17*^*EGFP*^ and *flt1_9a:nls-cherry* are co-expressed in individual LECs of the initial lymphatic arc. p’ shows high magnification of the domain of expression (blue dashed ROI). q-t, *sox17* overexpression results in loss of lymphatic marker expression. q-r, Stage IV AF (demarcated by yellow dashed lines) showing expression of *lyve1b:dsRed*, at the dorsal end of the AF (q), which is lost following mosaic expression of *UAS sox17* in *hsp70l:Gal4* animals (r, asterix). s,t, Confocal images of a 4.5 dpf *Tg(fli1a:Gal4; lyve1b:dsRed)* embryo showing the TD (s, arrowheads) that is lost (t, asterix) following mosaic expression of *UAS sox17*. Scale bars, 10µm (h), 15µm (b,k’), 20µm (b’,s,t), 30µm (a-a’’,c,f,k,m,p,p’), 50µm (d,l,n,o), 100µm (e,q,r), 200µm (i), 300µm (g). TD, Thoracic duct; DA, Dorsal Aorta; PCV, Posterior cardinal vein; ECs, Endothelial cells; LECs, Lymphatic endothelial cells.

Multiple lines of evidence confirmed the lymphatic identity of these vessels. First they are directly connected to the trunk lymphatics (Fig. 1c,c’, Extended Data Fig. 2d-d’’; Supplementary Video 1); second, they are labeled by several reporters highlighting LECs, including *prox1a, mrc1a* (Fig. 1c-h), *Tg(lyve1b:dsRed)*^15^ (Fig. 1i-k) and *Tg(flt4:mCitrine)*^16^ (Extended Data Fig. 2e-g’). In addition, we confirmed the specificity of the *prox1* transgene using single molecule fluorescent *in situ* hybridization (smFISH) on whole mount AFs. As seen in Fig. 1m,n, we detected clear expression of *prox1* mRNA both in the initial lymphatic arc (stage I), and in the lymphatic sprouts (stage II) further validating the lymphatic nature of these vessels. Finally, the identity of lymphatic vs. blood vessels was functionally confirmed by intravascular injection of Qdot705, which remain within the constraint of the blood system^5^ (Extended Data Fig. 1e-e’) and were hence detected only in the *kdrl+* blood vessels, and not in the vessels characterized as lymphatics (Fig. 1o-o’). Altogether our anatomical, molecular and functional analyses indicate that the early development of the AF is uniquely correlated with lymphatic, rather than blood vessel growth.

Strikingly, we noticed that as AF growth proceeds towards its mature form, the expression of most lymphatic markers, besides *flt4:mCitrine*, is gradually lost along the fin rays and becomes restricted to the dorsal most part of the AF, just ventral to the endo-exo skeletal junction (Fig. 1p-q, Extended Data Fig. 2h-i’). The disappearance of LEC markers was not due to vessel pruning or retraction, as all fin vessels could still be detected in the pan-endothelial *Tg(fli1a:dsRed)*^17^ (Fig. 1r). Moreover, the pattern and distribution of lymphatic markers detected at stage IV was maintained in AF vessels through adulthood, indicating this to be its final form (Fig. 1s-s’’). We also ruled out the possibility that the loss of lymphatic markers is related to the maturation of lymphatic vessels, as all transgenic reporters were readily detected in lymphatic vessels of other organs in the adult fish, including the heart^5^, meninges^18^ and the skin (data not shown).

The above results prompted us to investigate whether the mature AF blood vasculature in fact derives from the initial lymphatic vessels, via trans-differentiation of LECs. Notably, while lymphatic vessels have been shown to originate from multiple cell sources^19–26^, LECs themselves have not been shown to serve as progenitors for any other cell types. In order to uncover the origins of the mature vessels of the fin rays, we developed an EC-specific multicolor lineage tracing. Specifically, we generated fish expressing the *brainbow* cassette^27^ under pan-endothelial *fli1a*^28^ regulation (Extended Data Fig. 3a) (to be referred as ‘*flibow*’ hereafter). Mating these fish with *Tg(hsp70l:CreER*^*T2*^) animals, rendered progenies enabling heat-mediated CreER^T2^ expression, and 4-hydroxytamoxifen (4-HT) mediated CreER^T2^ activation in an ubiquitous manner^29,30^, allowing EC-specific *fli1a*-driven expression of different fluorophores (Extended Data Fig. 3b). We used confocal imaging to obtain lambda stacks, to acquire the entire emission spectra of all fluorophores upon simultaneous excitation, thereby generating a ‘spectral signature’ of individual ECs (Extended Data Fig. 3c,c’). While the heat and 4-HT treated progenies displayed ECs with diverse ‘spectral signatures’, siblings exposed to either treatment alone, contained only tdTomato-expressing ECs (Extended Data Fig. 3d-e’). In order to obtain highly efficient labelling of trunk lymphatics, we induced recombination at 60 hpf, before the onset of TD formation. Fluorescence from the ‘switched’ cells was readily detectable ∼24 hours post treatment (Fig. 2a), and it stabilized in the next 2 days along with a slight decay in the levels of pre-switched dTomato, as confirmed through reiterative imaging of the same animal (Fig. 2a-a’’ and Extended Data Fig. 3f,f’). Once stabilized, these signatures remained unchanged in ECs and their progenies for the entire experimental period (representative example in Extended Data Fig. 3g,g’). Following activation of the labelling, we screened *larvae* at 5-10 dpf (well before the initiation of AF formation) and selected those with differentially labeled trunk lymphatics (such as Fig. 2a’’). We then imaged each of these animals separately at different intervals to trace the origins of the initial AF plexus and the final AF vasculature. As seen in Fig. 2b-b’, we were able to detect distinctly labelled TD (Fig. 2b, green) and sprouts of the same origins in stage I AF (Fig. 2b’, green). These formed the Stage II plexus (Fig. 2c) and grew along the bones (Fig. 2d,e). Since in this fish only the anterior sprout was labelled (Fig. 2c, arrowheads) and not the posterior (Fig. 2c, arrow), the labelled LECs populated the anterior part of the AF (Fig. 2e). To our surprise, when tracing was carried out until the adult stage (3 mpf), we discovered that the entire AF blood vasculature in the adult fish, including the mid ray arteries and the peripheral veins, is in fact derived from a few LECs sprouting into the stage I AF (Fig. 2f,g, blue). In addition, imaging of animals with distinctly labeled blood vessel derived sprouts (Fig. 2h-h’’, BEC sprout-Green, arrowhead; LEC sprout – blue, arrow) confirmed our previous findings, namely that they remain restricted to the dorso-anterior region (Fig. 2i, arrowheads) without contributing to the mature fin ray vasculature (Fig. 2i’, arrows). Taken together, these results confirm that *bonafide* LECs in the early AF lose their lymphatic identity to become the blood vessels of the mature fin rays.

**Fig. 3:**
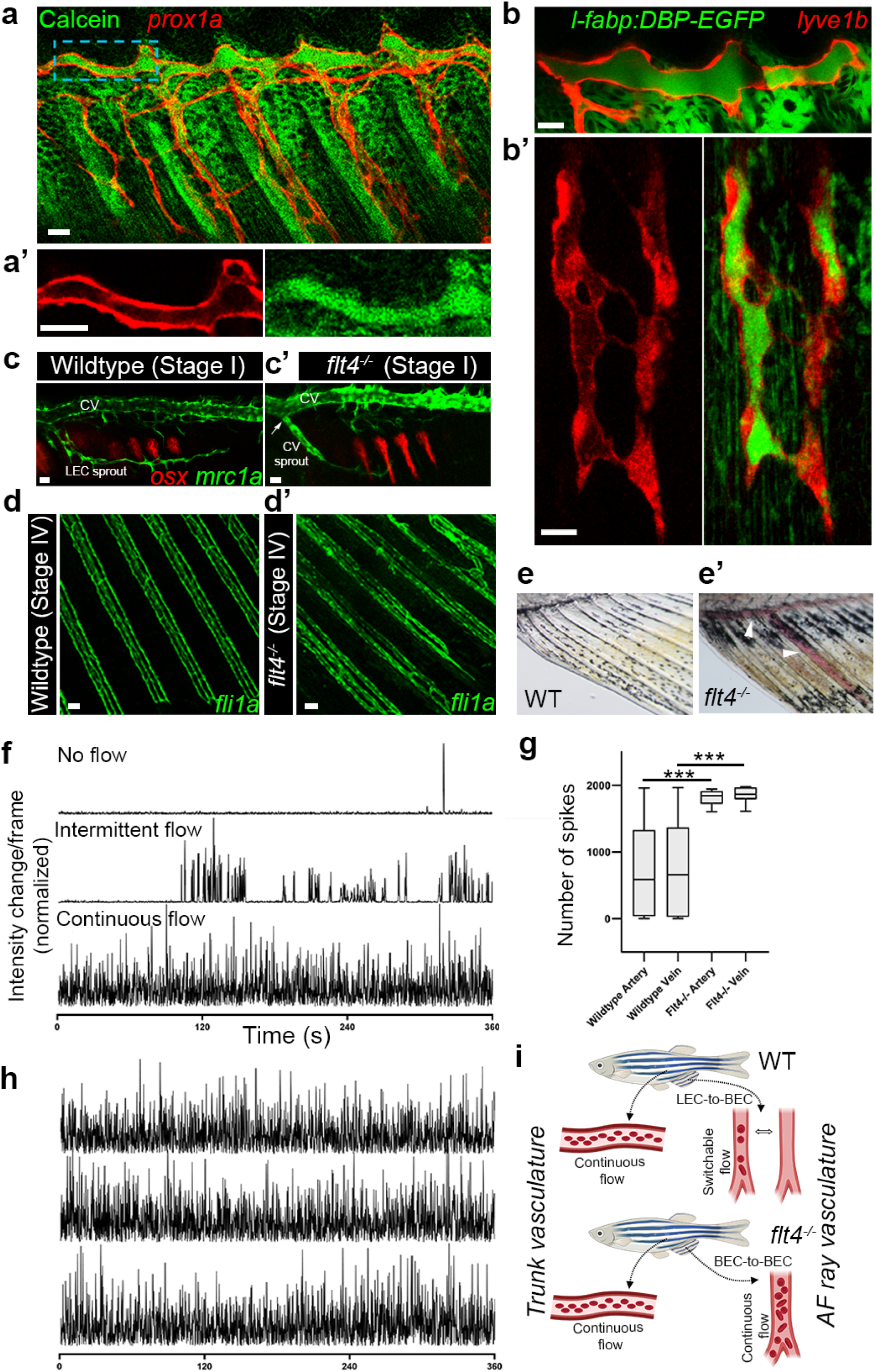
LEC-vs. BEC-derived blood vessels display different functional properties. a-a’, Distribution of calcein in stage III AF (a), following intramuscular injection in the trunk. a’ shows a close-up of a *prox1a+* lymphatic lumen (blue box in a) containing calcein. b-b’ *in vivo* imaging of the plasma tracer *l-fabp:DBP-EGFP* in stage II AF, depicting enrichment of plasma in the lumen of lymphatics (*lyve1b*), both in the main vessel (b) and the ventral sprouts (b’). c-d’, *flt4*^*-/-*^ animals vascularize their AF through blood vessel sprouts arising in the CV. Stage I AF lymphatic arc in WT (c), as compared to *flt4*^*-/-*^ where the sprout is directly connected to *mrc1a+* CV (c’, arrow). *osx:cherry* labelled osteoblasts are shown for morphological reference. d-d’, *fli1a:EGFP+* lumenized vessels are detected in stage IV AFs of both WT (d) and *flt4*^*-/-*^ mutant (d’) fish. e-e’, *flt4*^*-/-*^ displays erythrocyte pooling (e’, arrowheads) in the mature AF never detected in WT siblings (e). f-h, Differential erythrocyte flow in WT vs. *flt4*^*-/-*^ animals. Bright light intensity profile from the lumen of fin ray vessels in WT (f) or *flt4*^*-/-*^ (h) fish are shown, along with the quantification of erythrocyte-mediated intensity change (spikes) events (g). f,h, Intensity profiles from three representative animals, showing different modes of erythrocyte flow in wt (no flow, intermittent flow, continuous flow) (f), vs. continuous erythrocyte flow in all *flt4*^*-/-*^ mutants (h). i, Schematic illustration of the different flow properties in vessels originating via LEC-toBEC transition vs. BEC-to-BEC sprouting angiogenesis. Scale bars, 50 µm (d,d’), 20 µm (a,a’,b), 10µm (b’). ***, p<0.001. CV, Cardinal vein; BECs, Blood endothelial cells; LECs, Lymphatic endothelial cells.

Although the lymphatic EC fate has been shown to be “reprogrammable” under certain conditions presented by mutants or pathologies^31–34^, there is no evidence thus far of an *in vivo* physiological program that utilizes differentiated LECs to generate blood vessels. To investigate the molecular nature of this fate-switch, we analyzed previously published RNA-seq data derived from PROX1 downregulation in Human Dermal Lymphatic Endothelial Cells (HDLECs)^35^. In this setup, loss of lymphatic fate correlated with significant upregulation of a number of well-known blood vessel transcripts such as *KDR, FLT1, NR2F2, DLL4* and *EPHB2* (Fig. 2j, Extended Data Fig. 4a), as well as *BST1* (*CD157*) that has been shown to mark ECs displaying stem cell like properties^36^, and the transcription factor SOX17. Taking cue from this dataset, we asked whether some of these features are also part of the LEC-to-BEC transition taking place in the AF. We had previously shown that early lymphatic vessels in zebrafish originate from a population of PCV-resident angioblasts labeled by a *flt1*-enhancer-*Tg(flt1_9a_cfos:GFP)*^10^, whose expression is also enriched in arteries^5,37^, but undetectable in differentiated LECs (Extended Data Fig. 4b-b’). Unexpectedly, we noticed that around Stage II of AF formation, few LECs within the initial lymphatic arc turn on *flt1_9a:GFP* expression (Fig. 2k-k’’). As the rays form, additional *lyve1b* (Fig. 2l,l’) */prox1a* (Extended Data Fig. 4c) positive LECs begin expressing *flt1_9a:GFP*, regardless of whether they are part of the prospective arteries or veins, suggesting the progressive transition of these LECs toward a less differentiated state, reminiscent of their embryonic origins (Extended data 4d and ^10^). We then assessed the expression of Sox17, a transcription factor known to be enriched in mammalian arteries^38^, which was also upregulated in *PROX1*-deficient HDLECs (Fig. 2j, Extended Data Fig. 4a). Confocal imaging of a *Tg(sox17:EGFP)* reporter generated by knocking in an *EGFP* cassette into the *sox17* locus revealed that while *sox17*-directed expression is not detected in the initial LEC sprouts penetrating the AF (Extended Data Fig. 4e-f’), it becomes progressively enriched in the prospective fin ray arteries from the onset of stage II and onwards (Fig. 2m-o’). Notably, we could confirm, that the first LECs to express *flt1_9a* were also *sox17+* (Fig. 2p-p’), and that these are the cells that first lose the expression of lymphatic markers.The *sox17+* LECs, gradually upregulate also the expression of the BEC specific marker *kdrl*, which is initially detected only in the rostral, *sox17-*negative, BEC-derived blood vessels (Extended Data Fig. 4g-g’, cyan). However, as the transition proceeds, *kdrl* becomes enriched in *sox17+* LEC-derived blood vessels as well (Extended Data Fig. 4h-h’, cyan). Although Sox17 is considered an arterial specific marker^38^, and as such labels the prospective ray arteries (Fig. 2o-o’, arrowheads), we could also spot its expression in the veins of stage III AF (Fig. 2o-o’, arrows), suggesting the possible involvement of this transcription factor in all LEC-to-BEC transitions. To further investigate this hypothesis, we injected *UAS:Sox17* DNA into one cell stage *Tg(hsp70l:Gal4*^39^; *lyve1:dsRed)* embryos. Heat activation at 3 wpf rendered 25.24±8% of the injected fish (N=2, n=65) displaying patches of missing *lyve1* expression in the AF that was never observed in heat-shocked uninjected siblings (Fig. 2q,r, Extended Data Fig. 4i-j’’). Moreover, mosaic overexpression of Sox17 in ECs through injection of *UAS:Sox17* DNA into one cell stage *Tg(fli1a:Gal4*^40^; *lyve1b:dsRed)* embryos resulted in absence (Fig. 2s,t) or incomplete formation (Extended Data Fig. 4k-l’) of the TD in 17.94% of the injected embryos (N=2, n=234), as compared to 0 (N=2, n=87) of uninjected siblings. Taken together, our results reveal that certain LECs in the AF plexus gradually turn on blood vessel specific programs, while downregulating lymphatic marker expression and highlight Sox17 as a novel suppressor of LEC fate.

**Fig. 4:**
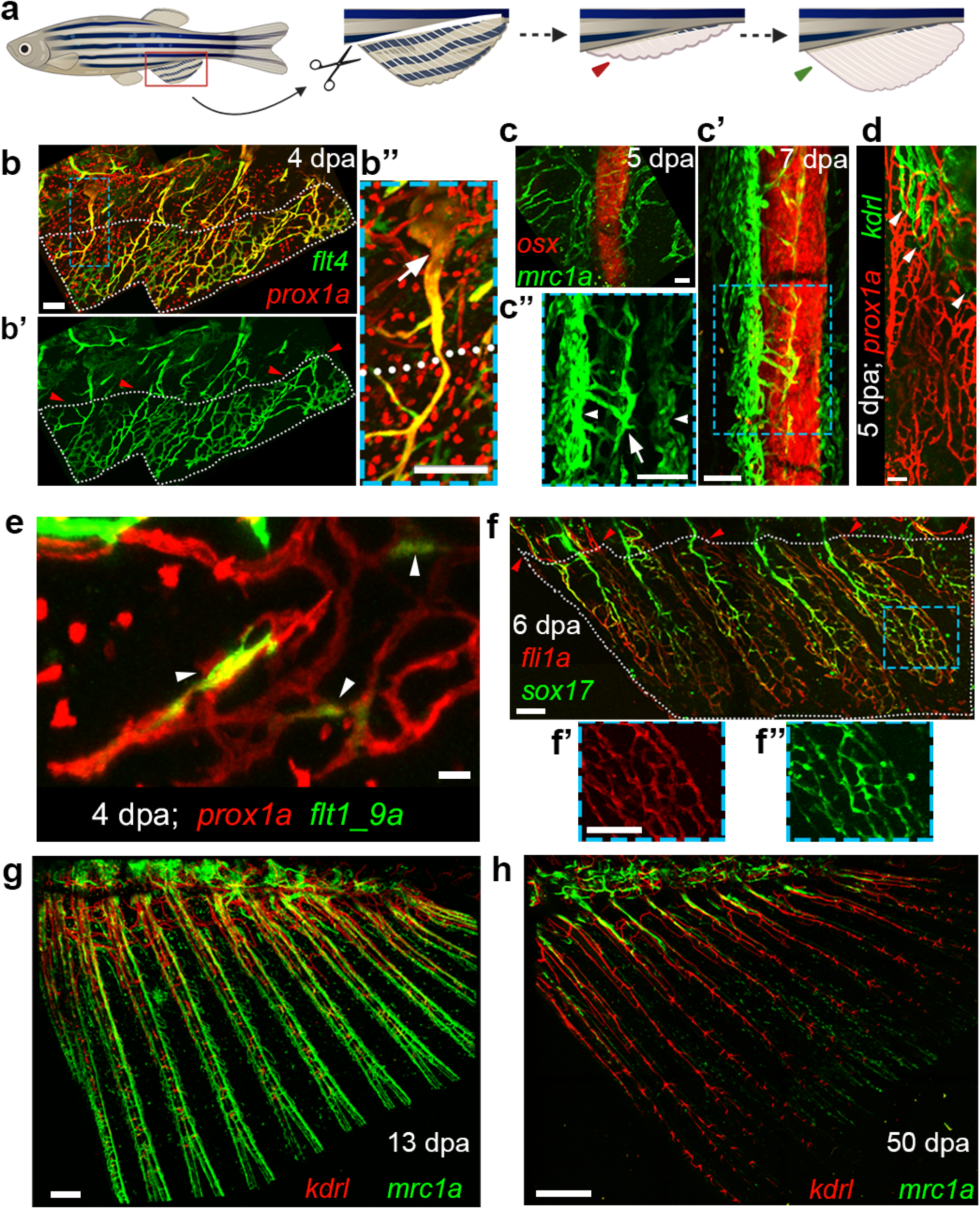
Regenerating AF are vascularized through LEC-to-BEC transdifferentiation. a, Schematic representation of AF regeneration protocol. The AF is amputated at the level of the trunk-fin junction, resulting in formation of a blastema (red arrowhead) from which the entire fin is reestablished (green arrowhead). b-b’’, Expression of *prox1a:RFP*;*flt4:mCitrine* (b,b’’) and of *flt4* only (b’) in the ECs re-vascularizing the regenerating AF (bounded by white dots) at 4 dpa (days post amputation). Pre-existing lymphatics (b’’, white arrow) from unamputated area contribute to the ECs of the regenerating tissue. c-c’’, *mrc1a:EGFP* labeled LECs growth along *osx:cherry* labeled bones as detected in 5dpa (c) and 7dpa (c’,c’’) AFs. Inset (c’’) shows LECs differentiating into mid-ray artery (arrow) and peripheral veins (arrowheads). d, Status of *prox1a:RFP+* lymphatics and *kdrl:EGFP+* blood vessels in a 5dpa regenerating AF, with arrowheads depicting the extent of blood vessel coverage in the tissue. e, *flt1_9a:GFP* expression is detected in *prox1a:RFP+* LECs of the regenerating AF plexus at 4dpa (arrowheads). f-f’’, Expression of *sox17*^*EGFP*^ in the 6dpa regenerated AF vasculature (highlighted by *fli1a:dsRed*). Insets (f’,f’’) show that the majority of ECs display *sox17*^*EGFP*^ expression. g-h, Status of *mrc1a:EGFP+* and *kdrl:cherry+* ECs depicting the differences between an early (g, 13 dpa) and a later (h, 50 dpa) time points of regeneration. In b-b’’,f the area of the regenerating AF indicated by red arrowheads represents the plane of amputation. In c-c’’,d and e the entire image is shown from a regenerated area, whereas for g and h, the entire regenerated fin is presented. Scale bars, 500 μm (h), 300 μm (g), 100 µm (b,b’’,c,c’’, f-f’’), 50 µm (d) and 20 µm (e).

During embryonic development, different cell populations have been shown to originate from multiple sources ^41,42^. However, the functional relevance of deriving a specific cell type from ontologically distinct progenitors, remains an open question. The case of AF vascularization is particularly interesting since, in spite of the presence of a conventional source of blood vessels (the CV) in the vicinity, the tissue utilizes an unusual source-lymphatic vessels. One important outcome of preferential lymphatic vascularization in the developing AF is that it creates a local hypoxic environment, largely regarded as key for bone formation and regeneration^43–46^. However, it has remained unclear how in such setup, devoid of blood vessels, do these rapidly developing structures receive the growth factors and minerals required to support bone growth. This seems particularly relevant in the developing AF because at the onset of fin ray growth, the associated endothelium is still ‘lymphatic’ (i.e. not connected to the blood circulation (Fig. 1o,o’)) and yet, osteoblast differentiation (Fig. 1f,g) and ray calcification occur, as highlighted by calcein staining of calcium deposits^8,47,48^ (Extended Data Fig. 5a). We thus hypothesized that in this case, recruitment of lymphatic vessels, provides a unique solution enabling delivery of necessary factors required for bone formation, while preserving hypoxic conditions. To test this hypothesis, we injected calcein intramuscularly in the anterior trunk, at 19dpf, with the AF at stage III onset. Within 10-15 mins, the calcium indicator was readily detected in the lumen of the AF lymphatics, as well as in the developing bones and associated mesenchyme (Fig. 3a,a’). Given that free calcium is primarily confined to plasma, we confirmed that in spite of not being connected to blood circulation, stage I AF lymphatic vessels do carry plasma, as seen in *Tg(l-fabp:DBP-EGFP)*^49^ animals (Fig. 3b,b’ and Extended Data Fig. 5b,b’). Together, these results strongly support the existence of a path for delivery of solutes into the developing AF, through the initial lymphatic plexus that concomitantly allows a hypoxic microenvironment.

As AF growth proceeds and LECs continue trans-differentiating, the newly generated, *sox17+*, LEC-derived blood vessels grow dorsally (Extended Data Fig. 5c, arrow) and establish patent connection to the dorsal aorta (DA) (Extended Data Fig. 5d-d’’). Consequently, stage III AF vessels are fully perfused by intravascularly injected Qdots705 (Extended Data Fig. 5e-e’) as opposed to stage II vessels (Fig. 1.o,o’), suggesting this to be the stage when they start functioning as blood vessels. Next, we asked whether the functionality of these vessels depends on their peculiar origins (lymphatic vessels) as opposed to a more conventional, blood vessel source. To answer this question we turned to *flt4*^*um203*^ animals^5,50^ that harbor a mutation in the Vegfc receptor, *flt4*, and are therefore devoid of trunk lymphatics (Extended Data Fig. 6a,a’). Interestingly, in the absence of lymphatic-derived sprouts, blood vessels arising from the overlying CV vascularize the AF (Fig. 3c-c’). Even though the AF vasculature shows slightly altered patterning in *flt4* mutants, the main vascular components are present along the rays in the adult fish (Fig. 3d,d’), providing an opportunity to compare blood vessels in the same organ arising from two different sources. The first thing we noticed is that *flt4* mutant AFs presented unusual blood pooling and accumulation of erythrocyte clusters, that were never detected in wt siblings (Fig. 3e,e’). To understand this phenomenon we recorded erythrocyte flow in the AF vessels of adult wt and *flt4*^*-/-*^ animals and found that mature wt AFs display an unusual pattern of ‘intermittent’ flow, with the majority of the fin rays completely lacking erythrocytes, while some switched between states of flow and stasis (Fig. 3f). This was even more dramatic in stage IV juvenile AFs, where we could not detect erythrocyte flow at all (Extended Data Fig. 6b). Thus, in spite of being the lymphatic-derived blood vessels of the mature AF, fully lumenized and perfused (Extended Data Fig. 6e,e’), they are devoid of erythrocytes. In contrast to this scenario, the AF vessels of adult *flt4*^*-/-*^ animals (which formed through angiogenic sprouting from the CV), display continuous blood flow and contain high erythrocyte density both in juvenile (Extended Data Fig. 6c,d, Supplementary Video 2) and adult fish (Fig. 3g,h, Supplementary Video 3). Thus, the source of the ECs (blood vs. lymphatic) forming the AF blood vasculature, determines the functional properties of these vessels in the mature fin, highlighting the importance of cell ontogeny for tissue functionality (Fig. 3i).

Thus far, we demonstrated how the phenomenon of LEC-to-BEC transdifferentiation is utilized to support AF growth during metamorphosis, a unique phase in the fish life-cycle characterized by multiple physiological changes^51^. We thus wondered whether similar mechanisms underlie revascularization in response to tissue damage and regeneration in adult animals where metamorphosis-specific changes are not relevant. To answer this question, we modified well-established protocols triggering caudal fin regeneration following amputation^52^, to surgically remove most of the AF in mature fish (>3mpf), leaving very little preexisting rays and associated vessels (Fig. 4a). As these animals regenerated their fins, the newly forming vessels featured renewed expression of lymphatic markers (Fig. 4b-d), which, as shown before, are no longer expressed by mature AF ECs (Fig. 1s-s’’). The regenerating AFs received sprouts originating from pre-existing lymphatics from the unamputated AF-trunk junction (Fig. 4b,b’’). Similar to the initial development of AF vessels in juvenile fish, these LECs grew in association with the regenerating bones, first as an ill-organized plexus (Fig. 4c) and later on establishing the more organized three vessel pattern (Fig. 4c’,c’’). Also here, the lymphatic coverage was more extensive than *kdrl+* blood vessels (Fig. 4d). The lymphatic identity of these vessels was progressively lost as shown by the presence of LECs expressing *flt1_9a:GFP* (Fig. 4e) and *sox17* (Fig. 4f-f’’), indicating their transitioning to a BEC fate. Finally, as the entire fin is fully regenerated, akin to stage IV of tissue morphogenesis (Fig. 1p-r), the lymphatic markers are lost from these vessels (Fig. 4g, h). These results indicate that adult AF regeneration brings about a similar program of LEC-to-BEC transition as detected during metamorphosis, reinforcing the importance of this mechanism for generating functional bone-associated vasculature of the fin. The fact that the regenerating AF utilizes this mechanism, in spite of the presence of a fully differentiated blood vascular network, indicates that LECs harbor the plasticity to give rise to blood vessels throughout life, and emphasizes the requirement for this specific EC origin.

Overall, our work highlights a new mechanism of blood vessel formation through LEC trans-differentiation, and provides the first *in vivo* evidence for a link between cell ontogeny and functionality in ECs. In conjunction with the growing body of literature revealing newer roles for lymphatic vessels^46,53– 57^, our findings shed light on our current understanding of lymphatic plasticity, diversification and their physiological consequences.

## Acknowledgements

The authors would like to thank Hila Raviv, Lital Shen and Dean Robinson (Weizmann Institute, Israel) for technical assistance, Dr. Kamalesh Kumari for assistance with image analysis and illustrations, Gabriella Almog, Roy Hofi, Alla Glozman and Anna Tatarin for superb animal care. The authors are grateful to all the members of the Yaniv lab for discussion, technical assistance and continuous support. This work was supported in part by European Research Council (818858) to KY, Israel Science Foundation (861/2013) to KY, Binational Science Foundation (2015289) to KY, Minerva Foundation (712610) to KY, the H&M Kimmel Inst. for Stem Cell Research, the Estate of Emile Mimran (SABRA program). K.Y. is the incumbent of the Enid Barden and Aaron J. Jade Professorial Chair and is supported by the Willner Family Center for Vascular Biology; the estate of Paul Ourieff; the Carolito Stiftung; Lois Rosen, Los Angeles, CA; Edith Frumin; the Fondazione Henry Krenter; the Wallach Hanna & Georges Lustgarten Fund, the Polen Charitable Trust and the Daniel Shapiro Cardiovascular Fund. R.N.D. was supported by EMBO long-term fellowship (ALTF 1532-2015), Edith and Edward F. Anixter Postdoctoral Fellowship and a senior postdoctoral fellowship by the Weizmann Institute of Science. K.D.P. acknowledges support from National Institutes of Health (R35 HL150713, R01 AR076342). W.H. was supported by the Deutsche Forschungsgemeinschaft (HE4585/4-1) and by the North Rhine-Westphalia ‘‘return fellowship’’.

## Author contributions

R.N.D designed and conducted experiments, analysed data, and co-wrote the manuscript; I.B., N.M., G.L., and Y.T. conducted experiments and data analyses; S.S. and Y.T. performed bioinformatics analyses; Y.H. and K.D.P. generated knockin zebrafish and analyzed data; M.B and J.N. generated transgenic lines; D.H. and R. E.-H. assisted with smFISH experiments; W.H. and K.D.P. provided transgenic lines and reagents; K.Y. directed the study, designed experiments, analyzed data and co-wrote the paper with inputs from all authors.

## Materials and Methods

### Zebrafish husbandry and transgenic lines

Zebrafish were raised by standard methods^58^ and handled according to the guidelines of the Weizmann Institute Animal Care and Use Committee^28^. The *Tg(lyve1:dsRed)*^*nz101* 15^, *Tg(fli1:dsRED)*^*um13* 17^, *Tg(flt1_9a:EGFP)*^*wz2* 10^, *Tg(mrc1a:Egfp)*^*y251* 4^, *Tg(prox1aBAC:KalTA4-4xUAS-E1b:uncTagRFP)*^*nim* 10^, *flt4*^*um203/um203* 50^, *Tg(kdrl:EGFP)*^*s843* 13^, *Tg(kdrl:tagBFP)*^*mu293* 59^, *Tg(osx:mCherry)*^*zf131* 8^, *Tg(flt4:mCitrine)*^*hu7135* 16^, *Tg(hsp70l:Gal4)*^39^, *Tg(l-fabp:DBP-EGFP)*^49^, *Tg(kdrl:HsHRAS-mCherry)*^*s916* 60^ and *Tg(fli1:Gal4)*^*ubs3* 40^ were previously described.

Experiments were conducted on fish from the same clutch. Initially fish were selected based on their size (∼5.2 mm for Stage I initiation). Selected fishes were subsequently staged based on criteria described in Table 1, Extended Data Fig. 1f-i. For regeneration experiments, animals were selected by their age (3-6 mpf).

For imaging, we either used *casper* (*roy*^*-/-*^; *nacre*^*-/*-^) ^61^ background, or the embryos were treated for 6 days with 0.003% N-phenylthiourea (PTU) (Sigma, St Louis, MO) to inhibit pigment formation.

### Generation of new transgenic lines

The *Tg(fli1a:Brainbow1*.*0L)*^*mu254*^ *(flibow)* transgenic fish was generated by cloning the CMV-Brainbow-1.0 L construct (Addgene #18721) downstream of the *fli1a* promoter^62^, into a plasmid containing *tol2* sites. The *Tg(hsp70l:CreER*^*T2*^) line was established by fusing the coding sequence of *CreER*^*T2*^ downstream of the heat-inducible promoter – *hsp70l* ^63^. The *Tg(flt1_9a_cfos:nls-cherry)* reporter was established by cloning the previously described *flt1_9a* enhancer^10^ into a *cfos:nlsCherry* destination vector. All plasmids were generated using the Gateway system as previously described^62^.

### Generation of *sox17*^*EGFP*^ knock-in zebrafish

The *sox17*^*EGFP*^ knock-in line (allele pd305) was generated by TALEN-mediated targeted knock-in through non-homologous end joining. Briefly, an obligate heterodimeric TALEN pair (left arm: 5’-GATCAATAAGGATACGC-3’, right arm: 5’-CGGGACTGCTCATCTC-3’) was assembled as described^64^ to induce a double strand break at the start codon region of the *sox17* gene. The donor vector has the same structure as described^65^ except that the TALEN sequences were replaced with the *sox17* right arm and another left arm (5’-GTATACTACTGCGGCTAT-3’) upstream of the EGFP-polyA cassette. Vector-free knock-in line was generated by co-injection of the TALEN mRNAs, donor vector and *flp* mRNA at one-cell stage.

### *Flibow* induction and analysis

Induction of Cre recombinase in *flibow* embryos was carried out at 2.5 dpf (60hpf). First the embryos were pre-heated at 36°C for 15 mins for triggering *hsp70l* mediated Cre expression. Then they were subjected to combined heat (36°C) and 5μM 4-Hydroxytamoxifen (diluted in E3 medium) treatment for 45mins. Upon completion of the treatment, embryos were washed three times in fresh E3 medium and returned to fresh water. Animals were then imaged at the appropriate AF development stages days and returned back to fresh water until the endpoint of the experiment.

### Sox17 overexpression

#### DNA constructs

To prepare the *UAS-sox17* construct, the coding sequences of *sox17* (NM_131287.2) was amplified using the following primers:

*sox17* F: 5’ - atgagcagtcccgatgcg −3’

*sox17* R: 5’ -tcaagaattattatagccgcagt −3’

The resulting *sox17* coding sequence was then cloned downstream of UAS using Gateway cloning.

#### Overexpression

The *uas:sox17* construct was injected in *Tg(fli1a:Gal4;lyve1b:dsRed)* or *Tg(hsp70l:Gal4; lyve1:dsRed*) embryos at 1-cell stage. Heat shock was carried out at 37°C for 1h on 21dpf *Tg(hsp70l:Gal4; lyve1:dsRed*) animals (for juvenile induction).

#### Anal fin regeneration

Adult fish (between 3-6 mpf) were anesthetized by immersion into 0.04% tricaine (Sigma, St Louis, MO) and the AF were carefully detached using surgical blade and forceps. The animals were immediately allowed to recover in fresh water.

#### Angiography and lymphatic uptake assays

Microangiography was performed on anesthetized *larvae* and juvenile animals, by injecting Qtracker705 (Invitrogen Q21061MP) or Calcein (Sigma) in the heart using microinjection glass capillaries, as described^4^. In adults, heart was accessed in anesthetized animals by a small incision, followed by microinjection of Qtracker705^5^. In both cases, imaging was initiated within 5 mins of the injection and the animals were euthanized immediately afterwards.

Lymphatic uptake was assessed by intramuscular injection of Qtracker705, as described^66^.

#### Single molecule fluorescent *in situ* hybridization (smFISH)

The construction of the probe library, hybridization procedure and imaging conditions were previously described^67^. In brief, probe libraries were designed using the Stellaris FISH Probe Designer (Biosearch Technologies, Inc., Petaluma, CA). The *prox1a* library consisted of 48 probes, each of 20 bps length, complementary to the coding sequence of the gene (Table 2), and was coupled to Cy5 (GE Healthcare, PA25001). As a modification of the standard tissue smFISH protocol, whole mount zebrafish AFs were fixed in cold 4% PFA for 30 minutes and washed twice with PBS containing 0.3% Triton X-100 (Sigma) for 5 min, followed by cold 70% ethanol for 2.5 h. The samples were then washed in 2XSSC and incubated for 10min with Proteinase K solution at 37°c. The next steps were performed as described^67^. Formamide concentration of the wash and hybridization buffers was increased to 25%. Additionally, the pre-incubation with the wash buffer was extended to 90 minutes. Slides were mounted using ProLong Gold (Molecular Probes, P36934) and imaged on a Nikon-Ti2-E inverted fluorescence microscope with 100x oil-immersion objectives connected to iXon Ultra 888 CCD camera (Andor, Oxford Instruments), using Nikon NIS Advance Elements software.

#### Microscopy and imaging

Confocal imaging was performed using a Zeiss LSM780 or LSM880 upright confocal microscopes (Carl Zeiss, Jena, Germany) equipped with water immersed 20x NA 1.0 or 10x NA 0.5 objective lens. Euthanized animals were mounted using 1.5% w/v low melting agarose. For reiterative imaging of same animals a custom-built chamber was utilized^37^. z-stacks were acquired at 2–3 μm increments. Larger images were acquired using tile-scanning, and the images were stitched using Imaris Stitcher 9.3 or Zeiss’ Zen software.

*flibow* confocal imaging and lambda stack acquisition were performed by single, simultaneous scans (for each z-plane) with 458nm and 514nm single photon excitation lasers. Using Zeiss multichannel detector, the resulting emission spectra were collected into 11 channels, each detecting a range of 18nm from 454nm to 650nm, which encompassed emissions from all three fluorescent proteins of *flibow*.

Analysis of *flibow* images was performed by intensity measurement of each of the 11 channels for the selected ROIs (manually drawn to select single ECs). Normalized values of these intensities were plotted for different ECs for comparison. In certain cases, when ECs displayed similar intensity profiles and same origins, we averaged intensity values of these different ECs.

#### Image processing

Confocal images were processed off-line using Fiji^68^ version of ImageJ (NIH) or Imaris 9.3 (Bitplane). The images shown in this study are single-views, 2D-reconstructions, of collected z-series stacks. The colocalization channel was created using the Imaris ‘Colocalization Module’. Co-localization thresholds were set manually.

*flibow* images were processed using Imaris 9.3. Each of the 11 channels were given RGB values that corresponded to the wavelength collected. Since many channels collect the emission from only one fluorescent protein, we reduced all our images to show 8 of these same channels, leaving out three channels from the tdTomato emission (that is represented by 5 channels) that represents non-recombinant expression.

smFISH images were processed in Fiji. Average background signal was determined from ROIs from areas lacking any RNA puncta, and this was used for background subtraction from the entire image.

The 3D volume rendering was performed using Imaris 9.3 (Bitplane).

#### RNA-seq analysis

We used publicly available raw counts data of RNA-seq of cultured human dermal lymphatic endothelial cells (hDLECs) incubated with siPROX1 compared to untreated cells (control)^35^. Genes that were expressed in more than 2 samples, had at least 10 reads across all samples, and their mean expression was > 4, were selected for downstream analysis. R package DESeq2 (v1.26)^69^ was used to normalize raw counts, perform regularized log2 transformation (rlog) and to identify differentially expressed genes (DEGs) across samples (p-value<0.05). In addition, lfcShrink function using apeglm estimation was used for shrink log2 fold change (LFC) calculation, which allows to assess the expression changes in lowly expressed genes^70^.

### AF erythrocyte flow analysis

Three ROIs were selected spanning different AF rays (that included veins and artery) of each anesthetized juvenile or adult fish and 6 min time lapse images from a single z-plane were acquired. The imaging was performed using the transmitted light detector in the LSM880 confocal microscope, after manually determining the desirable contrast for each fin. The image size, zoom factor, pixel dwell time were kept constant for all experiments, allowing acquisition of images at a fixed frame interval of 0.152s.

The erythrocyte flux through the vessels were measured in Fiji. We plotted bright light intensity profile over time to reflect the state of erythrocyte flow. For this, a line ROI was drawn across the lumen the vessel. Erythrocytes crossing this ROI caused fluctuations in the bright light intensity measured through this ROI. We determined the threshold of minimum intensity change that corresponds to passing of a single erythrocyte, and the number of events above this threshold were counted as erythrocyte-mediated intensity spikes and quantified in our ‘spike count’ plots.

### Statistical Analyses

Statistical significance for three or more samples was calculated via one-way ANOVA followed by post hoc Tukey’s for multiple comparisons.

All numerical data are reported as mean values ± SEM and were analyzed using Prism 5 software (GraphPad Software, Incorporated, La Jolla, CA, USA).

