## Supplemental Files for "Generation of specialized blood vessels through transdifferentiation of lymphatic endothelial cells"

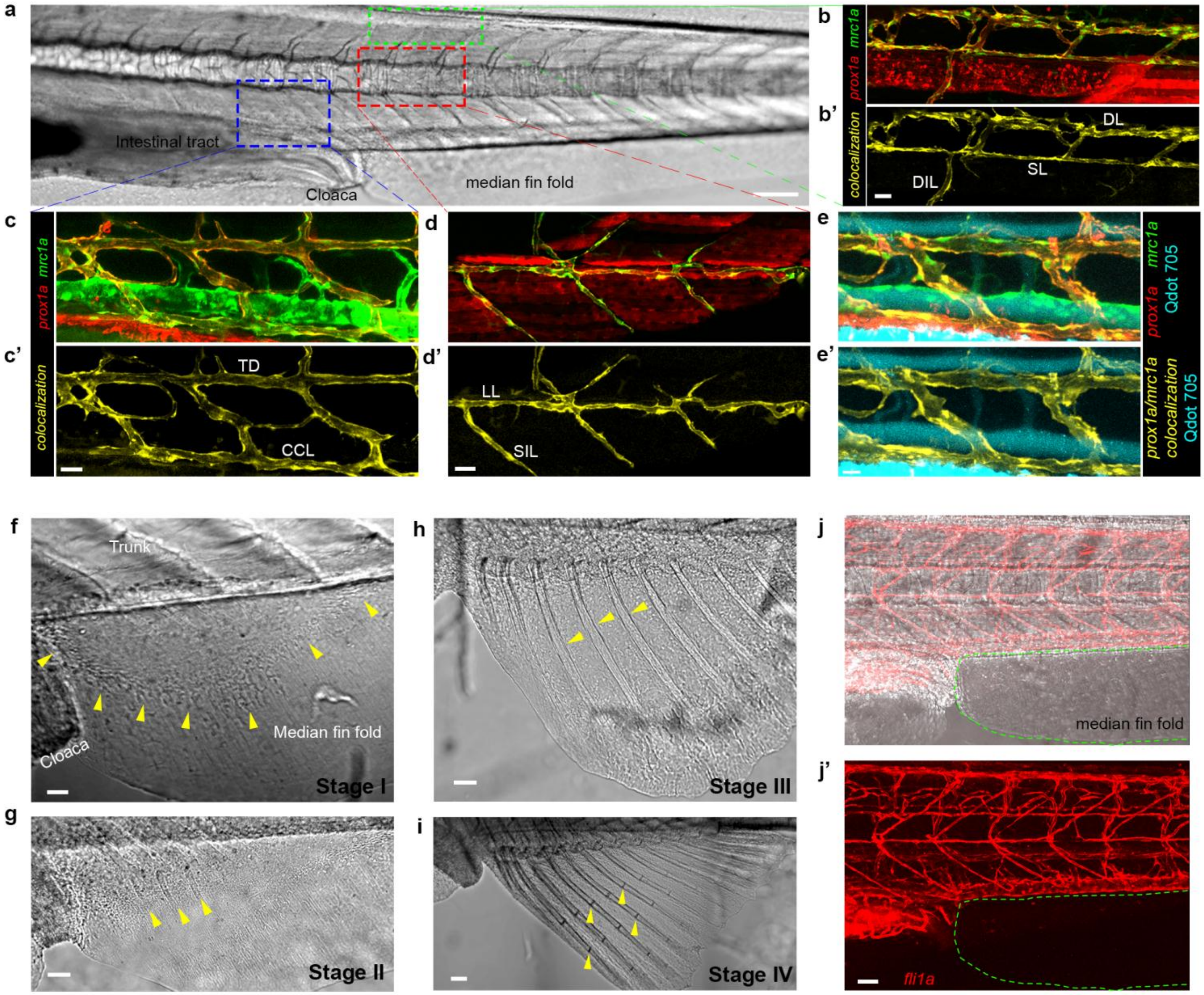


**Extended data 1: Stepwise formation of zebrafish anal fin.** a-d', *mrc1a:EGFP* and *prox1a:RFP* colocalization specifically highlights all lymphatic vessels in the larval trunk. a, Bright-light image of a 14dpf larval trunk, with boxes demarking dorsal (green), median (red) and ventral (blue) areas depicted in b-d. b,c,d, show complete *prox1a* and *mrc1a* transgene expression, b',c',d' depict colocalization channel. e-e', intravascularly injected Qdot705 (cyan) are detected in blood vessels but not in *mrc1a*/*prox1a* co-labelled lymphatics (colocalization highlighted in yellow). f-i, Bright-light images of AF development with arrowheads indicating important stage-specific characteristics, such as mesenchyme condensation (f), appearance of rays (g), ray growth (h) and formation of joints (i). j-j', ECs are not detected in the median fin fold (demarcated by green dashed lines) of *fli1a:dsRed larvae* prior to the initiation of AF development (Stage 0). Scale bars, 20µm (e'), 30µm (b',c',d',f), 50µm (g,h,j'), 100µm (i), 150µm (a). DL, Dorsal lymphatic, SL, Spinal lymphatic, DIL, Deep intersegmental lymphatic, TD, Thoracic duct, CCL, Collateral cardinal lymphatic, LL, Lateral lymphatic, SIL, Superficial intersegmental lymphatic.

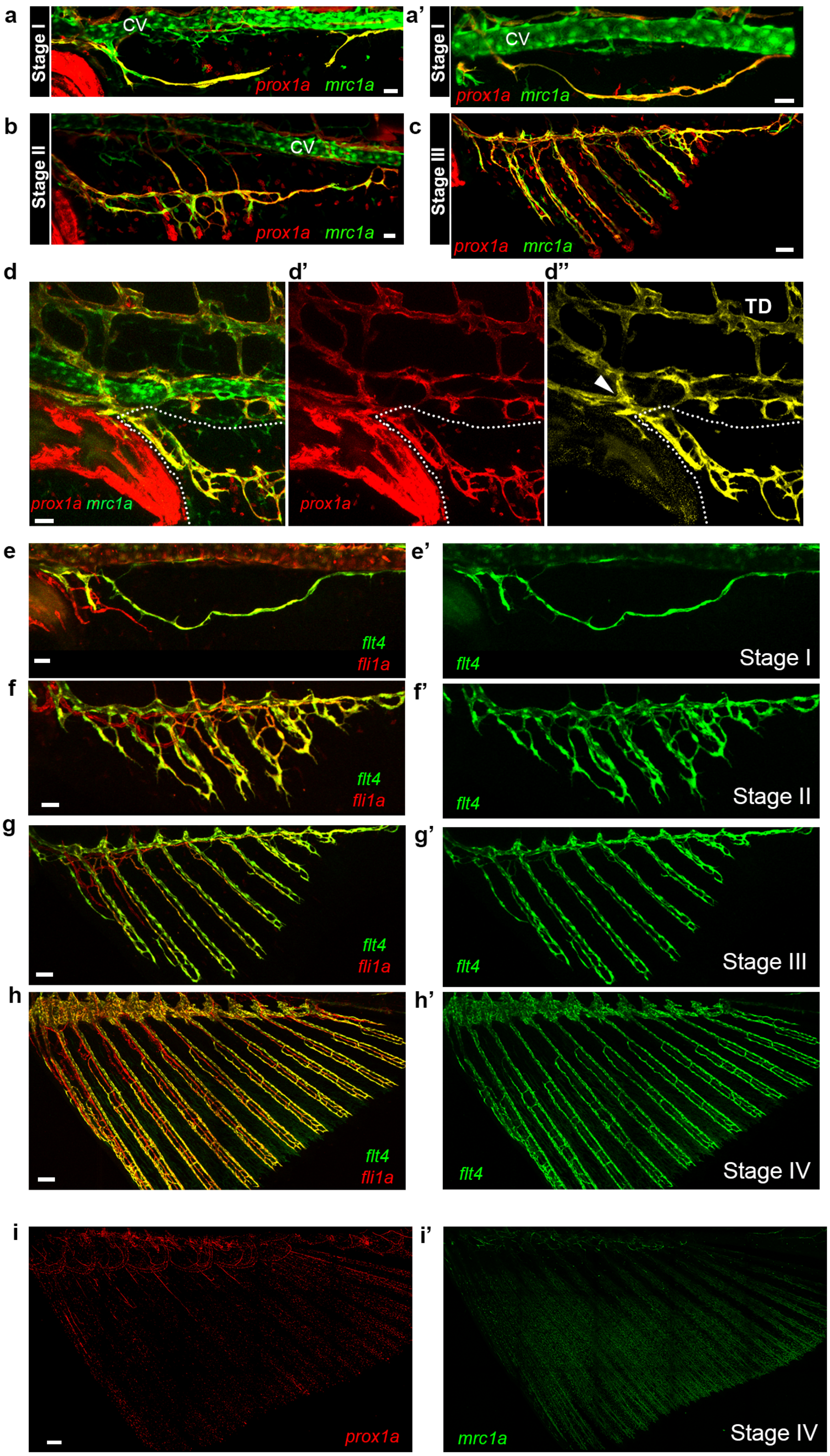
a
Stage I
CV
prox1a mrc1a
a'
Stage I
CV
prox1a mrc1a
b
Stage II
CV
prox1a mrc1a
c
Stage III
prox1a mrc1a
d
prox1a mrc1a
d'
prox1a
d''
TD
e
flt4
fli1a
e'
flt4
Stage I
f
flt4
fli1a
f'
flt4
Stage II
g
flt4
fli1a
g'
flt4
Stage III
h
flt4
fli1a
h'
flt4
Stage IV
i
prox1a
i'
mrc1a
Stage IV

**Extended data 2: Expression on lymphatic markers at different stages of AF morphogenesis.** a-c, Full expression of *prox1a:RFP* and *mrc1a:EGFP* colocalization channel depicted in Fig. 1c-e, showing lymphatics in stages I (a,a'),II (b) and III (c) of AF development. d-d'', AF lymphatics sprout from trunk lymphatics as shown in Tg(*mrc1a:GFP*;*prox1a:RFP) larvae*. White dots demarcate the developing AF tissue, and TD, thoracic duct. d'' shows colocalization channel, arrowhead points to the connection between the trunk and AF lymphatics. CV, Cardinal vein. e-g', Expression of *flt4:mCitrine* through stages I-III of AF development shown along with pan endothelial *fli1a:dsRed* (e,f,g) and separately (e',f',g'). h-h', Continued expression of *flt4:mCitrine* in the vessels of stage IV AF, shown with *fli1a:dsRed* (h) and separately (h'). i-i', Stage IV AF of *prox1a* (i) and *mrc1a* (i') fish, whose colocalization channel is depicted in Fig. 1q . Scale bars, 30µm (a,a',b,d,e,f), 50µm (c,g), 100µm (h), 200µm (i).

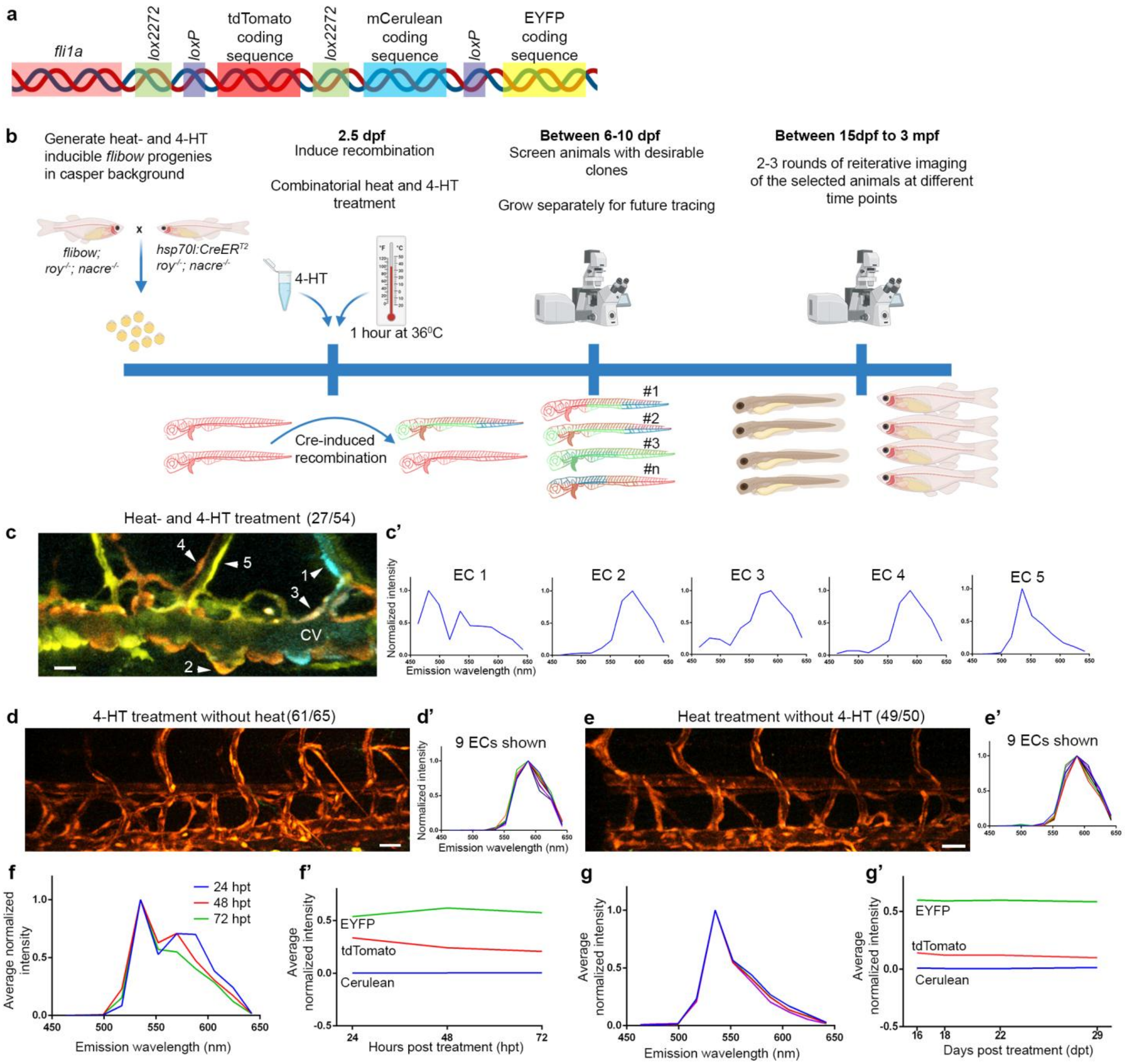


**Extended data 3: A platform for long-term EC lineage tracing.** a, Schematic representation of the *flibow* construct used to drive expression of *brainbow* under the *fli1a* promoter. Cre recombinase allows expression of different fluorescent proteins (*lox2272*- mCerulean or *loxP*-EYFP) in non-recombinant (tdTomato) animals. b, Schematic representation of the *flibow* lineage tracing protocol used in this study. c-e', Different representative examples of induced recombination. Combined heat- and 4-HT treatment allowed clear identification of 27 *larvae* (out of 54 screened) displaying differentially labelled ECs. Normalized intensity (c') plotted across a range of emitted wavelength (x-axis) from 5 CV-ECs (c, numbered arrowheads), depicting how ECs display a unique spectral signature, based on the levels of expression of the 3 fluorophores. 4-HT (d, 61/65) or heat treatment (e, 49/50) alone did not result in generation of recombinant outcomes. d’ and e’ depicts ‘spectral signature’ from 9 ECs from each group, demonstrating lack of diversity and presence of the tdTomato emission (non-recombinant outcome) signal only. f-g’, Assessment of the stability of the fluorophore ratios after Cre induction. e-e’, Average normalized intensity from a ‘switched’ clonal population (n=4) measured at 24, 48 and 72 hours post treatment (hpt), showing slight deviation with time (f). Intensities corresponding to each fluorophore are depicted separately for the 3 times-points and show a decay in tdTomato signal, that contributes to the deviation in the spectral signature shown in e. g-g’ Similar measurements carried out on ‘switched’ clonal ECs from another sample at 16, 18, 22, 29 days post treatment (dpt), shows a negligible deviation between the channels (f), along with stable intensity proportions between the fluorophores (f’). Scale bars, 30 μm (d,e), 15 μm (c). CV, Cardinal vein; ECs, endothelial cells.

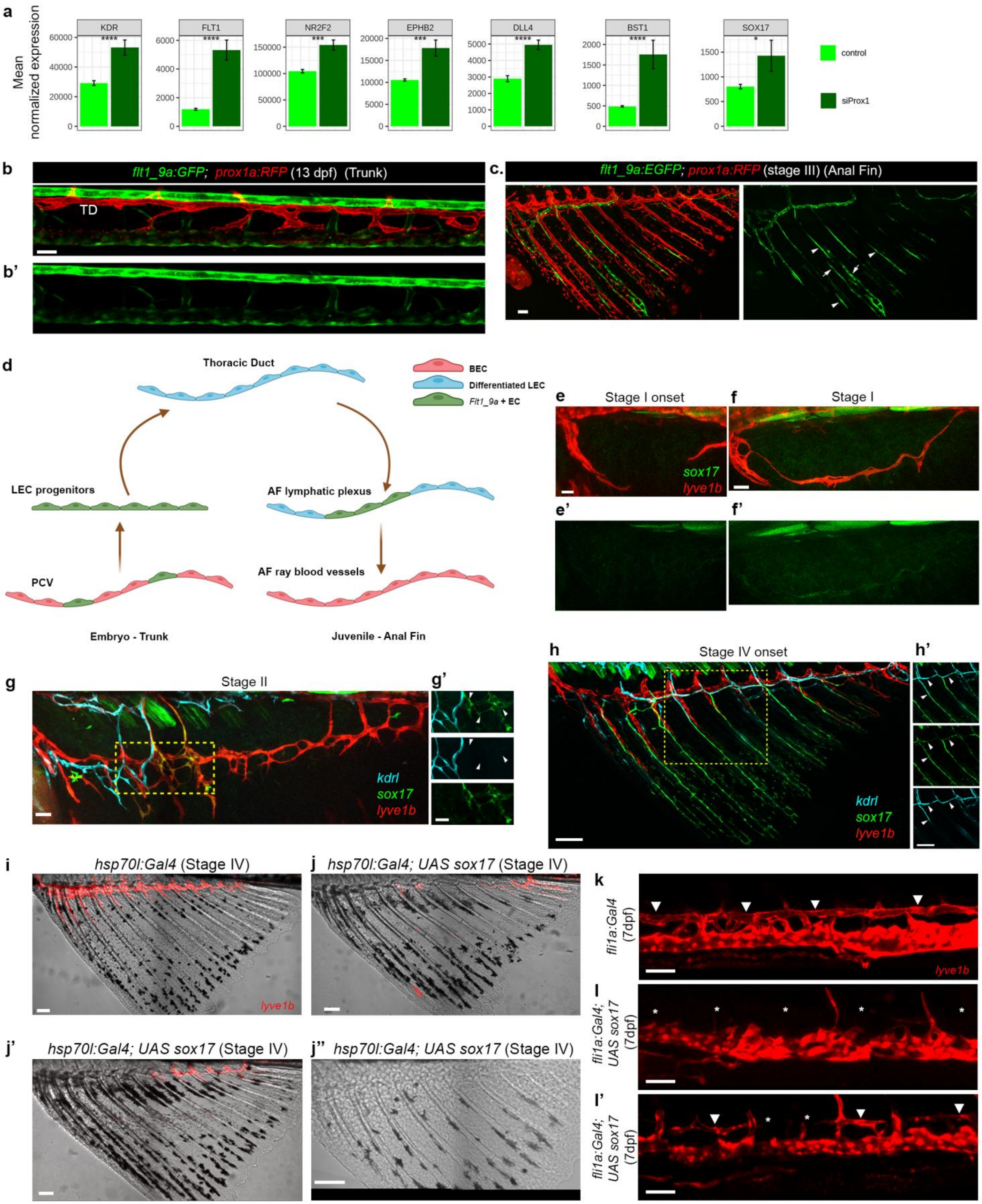
a
Mean normalized expression
KDR
****
FLT1
****
NR2F2
***
EPHB2
***
DLL4
****
BST1
****
SOX17
*
control
siProx1
b
flt1_9a:GFP; prox1a:RFP (13 dpf) (Trunk)
TD
b'
c.
flt1_9a:EGFP; prox1a:RFP (stage III) (Anal Fin)
d
Thoracic Duct
BEC
Differentiated LEC
Flt1_9a + EC
LEC progenitors
AF lymphatic plexus
PCV
AF ray blood vessels
Embryo - Trunk
Juvenile - Anal Fin
e
Stage I onset
f
Stage I
sox17
lyve1b
e'
f'
g
Stage II
g'
kdrl
sox17
lyve1b
h
Stage IV onset
h'
kdrl
sox17
lyve1b
i
hsp70l:Gal4 (Stage IV)
lyve1b
j
hsp70l:Gal4; UAS sox17 (Stage IV)
j'
hsp70l:Gal4; UAS sox17 (Stage IV)
j''
hsp70l:Gal4; UAS sox17 (Stage IV)
k
fli1a:Gal4 (7dpf)
lyve1b
l
fli1a:Gal4; UAS sox17 (7dpf)
l'
fli1a:Gal4; UAS sox17 (7dpf)

**Extended data 4: Loss of lymphatic fate is associated with gain of BEC molecular signatures.** a, Bar plots of normalized expression of selected genes that show upregulation following PROX1 suppression in HDLECs (siProx1) (****, p-value<0.0001; ***, p-value<0.001; **, p-value<0.01; *, p-value<0.05, Wald test). b-b’, *prox1a:RFP* labeled differentiated lymphatic vessels (TD) in 13 dpf larval trunk do not express *flt1_9a:EGFP*. c, Expression of *flt1_9a:EGFP* is shown along with *prox1a:RFP* + lymphatics in the prospective arteries (arrowheads) and veins (arrows) of stage III AFs. d, Schematic representation of the dynamics of *flt1_9a:GFP* expression during LEC progenitor specification in the early embryo and LEC-to-BEC transition in juvenile AF. e-f, *sox17:EGFP; lyve1:dsRed* stage I AFs showing no *sox17* expression in the sprouts (e-e’) and in the lymphatic arc (f-f’). g-h’, Gradual expression of *kdrl* is detected in s*ox17+* vessels of the AF. g-g’, LECs (*lyve1b:dsRed*), transdifferentiating LECs (*sox17:EGFP*) and blood vessels (*kdrl:BFP*) shown together (g) in stage II AF. Transdifferentiating LECs (g’, arrowheads) do not show expression of *kdrl* at this stage. h-h’, Same three labels in a different fish at the onset of stage IV (h), showing growing *sox17*+ vessels that start expressing *kdrl* (h’, arrowheads). i-l’, Mosaic overexpression of *uas:sox17* cause loss of *lyve1b:dsRed* expression with varying degrees of penetrance. i-j’’, Stage IV AF depicting normal expression of *lyve1b*, at the dorsal end of the AF in Tg(*hsp70l:Gal4; lyve1b:dsRed)* fish (i). Patchy (j,j’) or complete (j’’) loss of *lyve1b* is observed following *hsp70l:Gal4* mediated overexpression of *uas:sox17*. k-l’, Mosaic overexpression of *uas:sox17* in Tg(*fli1a:Gal4)* embryos, affects TD formation in 7dpf larvae (arrowheads point to TD, asterisk denote absent TD segments). Scale bars, 30µm (b,e,f,g,g’), 50µm (c,k,l,l’), 100µm (h,h’,i,j-j’’). TD, Thoracic duct.

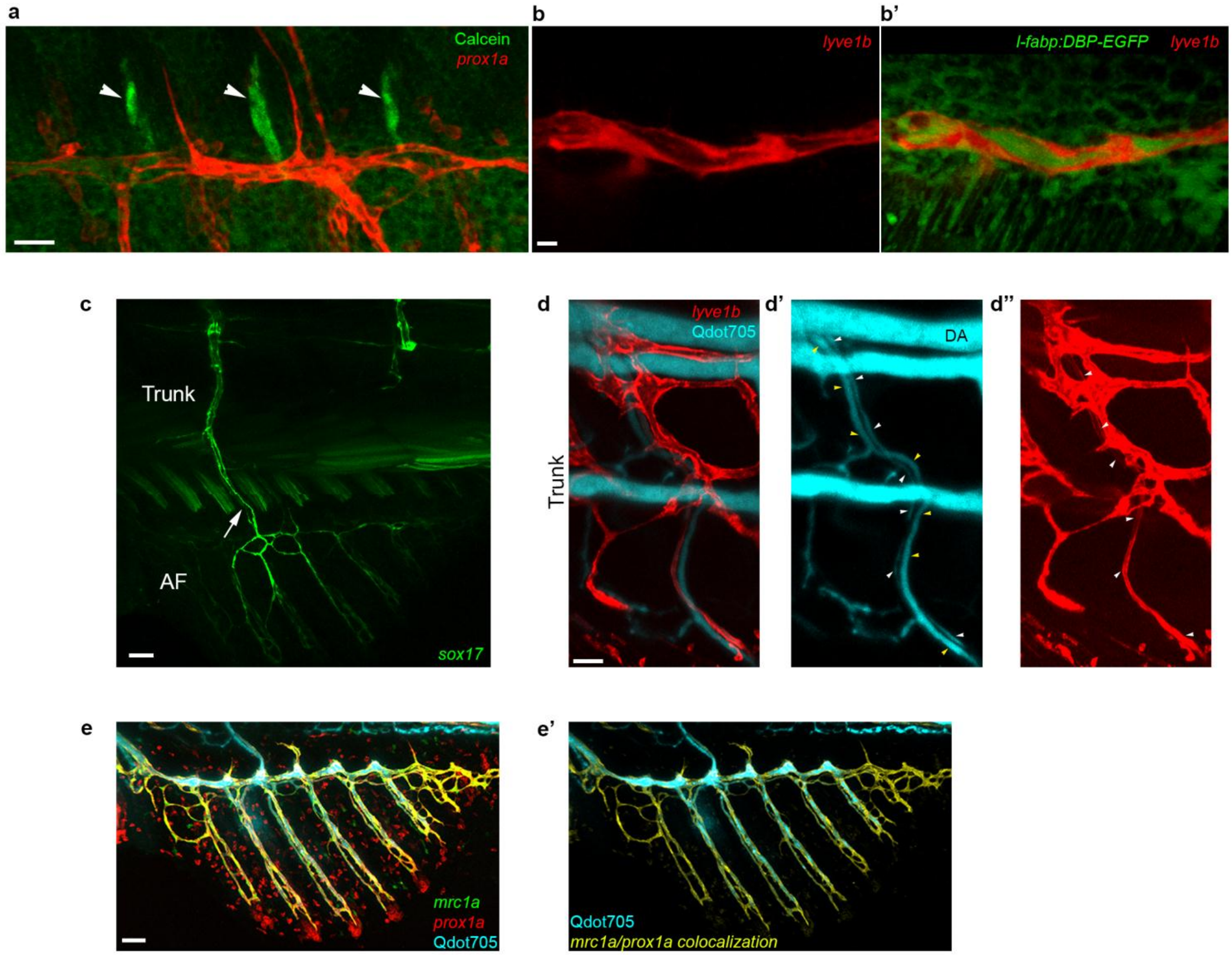


**Extended data 5: Functional characterization of the AF vasculature.** a, Calcified bones (arrowheads) in stage II AF, following intravascular injection of calcein. b-b', *lyve1b+* lymphatic arc in stage I AF (b), carrying circulating plasma, as visualized in the *l-fabp:DBD-EGFP* transgene (b'). c, *sox17+* vessels in stage III AF grow dorsally, towards the trunk vasculature. d-d'', Microangiography with Qdot705 depicts patent connections established between the DA and stage III AF (not seen in the image) through two vessels, one LEC-derived blood vessel (*lyve1+,* white arrowheads, d'-d'') and another lacking *lyve1* expression (yellow arrowheads, d'). e-e', Intravascular injection of Qdot705 (cyan) in the trunk of stage III AF fish results in labeling of *mrc1a*;*prox1a* positive AF vessels (e). e' shows *mrc1a*, *prox1a* colocalization channel along with Qdot705. Scale bars, 50 µm(c,e), 30µm (d), 20µm (a), 5µm (b).

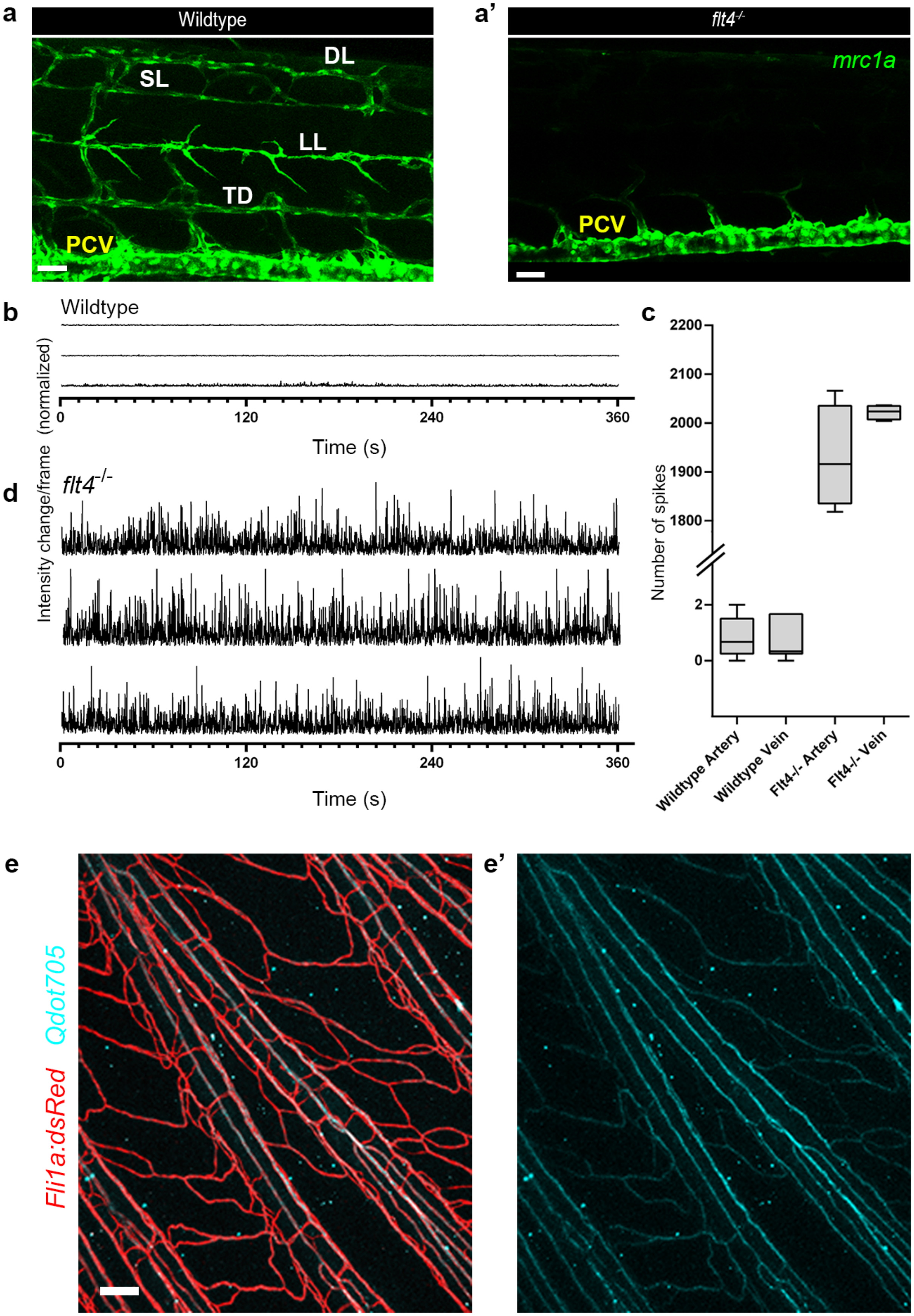

a
Wildtype
SL
DL
LL
TD
PCV
a'
flt4-/-
mrc1a
PCV
b
Wildtype
Intensity change/frame (normalized)
0
120
240
360
Time (s)
c
Number of spikes
2200
2100
2000
1900
1800
2
0
Wildtype Artery
Wildtype Vein
Flt4-/- Artery
Flt4-/- Vein
d
flt4-/-
0
120
240
360
Time (s)
e
Fli1a:dsRed
Qdot705
e'

**Extended data 6: Difference in erythrocyte flow properties in WT and *flt4*$^{-/-}$ juvenile zebrafish.** a-a', Trunk lymphatics (a) are absent in *flt4*$^{-/-}$ mutants (a'). b-d, Differences in erythrocyte flow in WT vs. *flt4*$^{-/-}$ juvenile fish (stage IV). Bright light intensity profile from the lumen of fin ray vessels in WT (b) vs *flt4*$^{-/-}$ (d) are shown, along with the quantification of erythrocyte-mediated intensity change (spikes) events (c). b,d Intensity profiles from three representative animals, showing no erythrocyte flow in WT (b), vs. continuous erythrocyte flow in *flt4*$^{-/-}$ mutants (d). e-e', Microangiography with Qdot705 (cyan) through the adult fish heart, showing presence of cyan labeling in *fli1a:dsRed* AF vessels (e) . Scale bars, 100 μm (e), 50 μm (a,a'). DL, Dorsal lymphatic , SL, Spinal lymphatic, LL, Lateral lymphatic, TD, Thoracic duct, PCV, Posterior cardinal vein.

**Supplementary Video 1:** 3D volume rendered image depicting the *mrc1a:EGFP* positive and *prox1a:RFP* negative CV (green) and the *mrc1a:EGFP*, *prox1a:RFP* double positive lymphatic vessels (yellow) that penetrate the Stage I AF. Scale bar, 50 μm.

**Supplementary Video 2:** Time-lapse video depicting erythrocyte flow along a fin ray in a wildtype (left, wt) and *flt4*$^{-/-}$ (right) stage IV AF. Yellow arrows point to the vessels. Images are shown for a 5min recording, with a frame interval of 0.152s and the video is being played at 50 fps.

**Supplementary Video 3:** Time-lapse video depicting erythrocyte flow along a fin ray in a wildtype (left, wt) and *flt4*$^{-/-}$ (right) adult fish AF. Yellow arrows point to the vessels. Images are shown for a 5min recording, with a frame interval of 0.152s and the video is being played at 50 fps.

**Table 1:** Stages of anal fin morphogenesis in the median fin fold, along with its morphological and endothelial features.

| Stage | Morphology | Endothelial cell pattern |
|---|---|---|
| **0** | Membranous fin fold (Extended Data Fig. 1j). | ECs completely absent. |
| **I** | Appearance of mesenchymal condensation (Extended Data Fig. 1f). | Trunk derived ECs sprouts along the edge of the condensation to form an arc, encircling the condensation. |
| **II** | Appearance of early fin rays, extending ventrally (Extended Data Fig. 1g). | ECs from the arc extend and proliferate ventrally to form a plexus. |
| **III** | Prominently visible fin rays without joints (Extended Data Fig. 1h). | The EC plexus gets organized along the rays, with two distinct vessels associated with each ray. |
| **IV** | Fin rays start to display joints and grow further ventrally (Extended Data Fig. 1i). | A third vessel is formed along the rays, and they resemble the three vessel (vein-artery-vein) unit associated with each fin ray in the adult fin. |

**Table 2 :** Table of *prox1a* smFISH probe libraries used in this study

| ***prox1a* smFISH probe sequences** | | | |
|---|---|---|---|
| 1. | tggaactgttattgtgccat | 25. | Ctctgtttggtcatttgagt |
| 2. | aagagggatgtgctgtcatg | 26. | Ccagtctgatctgaagagtt |
| 3. | tcctatgtcaactcttcttc | 27. | gagggaggactaaaacccat |
| 4. | tttgcgcgggtaaaaaccac | 28. | gaggaagggggaaaggatgg |
| 5. | tggggattcatggcactaag | 29. | acctaaggggctttggaaag |
| 6. | tgcaccactgaacactcaac | 30. | aggagaagacctatcctttc |
| 7. | ctcttcaacagcttgcgaag | 31. | ccgcattctgactttatgag |
| 8. | atcacagcgtcttcgtaaga | 32. | gatgtctgacatgtcctgaa |
| 9. | ctggaaattaggctcactgc | 33. | gcgcgtgtagaagaacatca |
| 10. | gctgaatggtgaaaggcact | 34. | acatcttcagcatgttggag |
| 11. | tccatctcaaattgggtcat | 35. | gcggttaaacttcacgtctg |
| 12. | cgttcttcatgtttctgttc | 36. | tgatcagctgggatgtgatg |
| 13. | ggtagaacttctcttggagt | 37. | aactcgcggaagttgctgaa |
| 14. | tcatcgttttcagagtcagt | 38. | ttgtagtgcatgttgagagc |
| 15. | aacgatctgggacagagtcg | 39. | tccggaacctcaaaatcgtt |
| 16. | aggtcagacatttcgttgtc | 40. | tggcattgaaaaactcccgt |
| 17. | gtgctcgatccaaaaagtgc | 41. | cttcttccaggaaggatcaa |
| 18. | tttagcgtttctgccaattg | 42. | aaaatctctgggacctcact |
| 19. | gacattgctgtattgagctc | 43. | agaagctcctgcagacaatt |
| 20. | aaaacctgaggaactgggcg | 44. | aggaatgctacctcttactt |
| 21. | tctccattgactgcgaaacg | 45. | ggggaaatggggtaaggatc |
| 22. | tggcagtatggaagtttgga | 46. | tcagttccatttgttaaccg |
| 23. | accaaaacattgcaagcgct | 47. | cattgcaaacatcctgggat |
| 24. | tccaggggattaggaatgat | 48. | ttattttctgtagcattgcc |
